# Voluntary Exercise Attenuates Tumor Growth in a Preclinical Model of Castration-Resistant Prostate Cancer

**DOI:** 10.1101/2024.10.16.617081

**Authors:** Nicolas Berger, Benjamin Kugler, Dong Han, Muqing Li, Paul Nguyen, Meaghan Anderson, Songqi Zhang, Changmeng Cai, Kai Zou

## Abstract

**Purpose:** To examine the effects of voluntary exercise training on tumor growth and explore the underlying intratumoral molecular pathways and processes responsible for the beneficial effects of VWR on tumor initiation and progression in a mouse model of Castration-Resistant Prostate Cancer (CRPC).

**Methods:** Male immunodeficient mice (SCID) were castrated and subcutaneously inoculated with human CWR-22RV1 cancer cells to construct CRPC xenograft model before randomly assigned to either voluntary wheel running (VWR) or sedentary (SED) group (n=6/group). After three weeks, tumor tissues were collected. Tumor size was measured and calculated. mRNA expression of markers of DNA replication, Androgen Receptor (AR) signaling, and mitochondrial dynamics was determined by RT-PCR. Protein expression of mitochondrial content and dynamics was determined by western blotting. Finally, RNA-sequencing analysis was performed in the tumor tissues.

**Results:** Voluntary wheel running resulted in smaller tumor volume at the initial stage and attenuated tumor progression throughout the time course (P < 0.05). The reduction of tumor volume in VWR group was coincided with lower mRNA expression of DNA replication markers (*MCM2*, *MCM6*, and *MCM7*), AR signaling (*ELOVL5* and *FKBP5*) and regulatory proteins of mitochondrial fission (Drp1 and Fis1) and fusion (MFN1 and OPA1) when compared to the SED group (P<0.05). More importantly, RNA sequencing data further revealed that pathways related to pathways related to angiogenesis, extracellular matrix formation and endothelial cell proliferation were downregulated.

**Conclusions:** Three weeks of VWR was effective in delaying tumor initiation and progression, which coincided with reduced transcription of DNA replication, AR signaling targets and mitochondrial dynamics. We further identified reduced molecular pathways/processes related to angiogenesis that may be responsible for the delayed tumor initiation and progression by VWR.

## INTRODUCTION

Prostate cancer (PCa) is the second most common cancer among men and the fifth leading cause of cancer related death worldwide (1). Androgen deprivation therapy (ADT) is often the first line of treatment in patients with PCa. Although most patients respond well to ADT in the initial stage, PCa cells eventually become resistant to this treatment, resulting in the development of castration-resistant prostate cancer (CRPC), leading to more aggressive cancer and lethality (2, 3). While current FDA-approved therapies such as enzalutamide (ENZ) and abiraterone (ABI) bring benefits to patients with CRPC, the frequent occurrence of drug resistance ultimately leads to the failure of therapy (4). Therefore, it is critical to develop effective interventions, including life-style changes, that prevent or slow down the progression of CRPC.

Mounting evidence from epidemiological studies have shown that exercise or increased physical activity reduced PCa incidence and delayed the progression to advanced PCa (5–9). For example, a prospective cohort study demonstrated that PCa patients who exercised for 3 or more hours per week had a significantly reduced rate of disease progression compared with those who exercised for less than an hour per week (5). These positive findings were further corroborated by intervention studies in preclinical animal studies using either voluntary wheel or involuntary treadmill running as an exercise regimen. However, to date, evidence of the effects of exercise on advanced PCa (i.e., CRPC) and the associated underlying mechanisms is scarce (10).

Recently, Yang et al. reported that three weeks of voluntary wheel running inhibited tumor growth in a syngeneic mouse model of CRPC by inhibiting cell proliferation and promoting apoptosis in tumors (11). However, the mechanisms by which exercise attenuates tumor progression remain not fully elucidated.

Mitochondrial alterations are common in PCa tumors, leading to increased mitochondrial metabolism to meet their energy requirement (12), which plays an important role in tumorigenesis and tumor progression (13–15). These organelles form dynamic networks that are capable of changing morphology through the regulation of mitochondrial dynamics processes (e.g., fusion and fission) (16). Mitochondrial fusion is regulated primarily by Mitofusins 1 and 2 (MFN1 and MFN2) and Optic atrophy protein 1 (OPA1) (17). Conversely, mitochondrial fission is primarily mediated by Dynamin-related protein 1 (Drp1), which can be activated and translocated to mitochondria to initiate fission process (17, 18). The orchestrated balance of the regulatory protein machinery among fusion and fission is vital in maintaining homeostasis of healthy cells (19). Dysregulated mitochondrial dynamics has been linked to dysregulated cancer cell metabolism and tumor microenvironment (20–22).

Aberrant expressions in regulatory proteins of mitochondrial dynamics machinery have been reported in PCa cell lines and tumor tissues, including upregulation of fission proteins, Drp1 and MFF, and fusion proteins, MFN11 and MFN2, in both androgen-sensitive and castration-resistant AR-driven PCa cell lines and PCa patient’s tumor tissues (15, 23, 24). Chronic exercise training is known to improve mitochondrial dynamics by reducing Drp1 activity and enhance mitochondrial function in various tissues (25–28). However, it is largely unknown how exercise alters mitochondrial dynamics regulatory machinery in tumor tissues.

The present study was sought to examine the effects of voluntary wheel running on tumor growth and the regulatory machinery of mitochondrial dynamics in a mouse model of CRPC. We hypothesized that voluntary wheel running would improve the regulatory machinery of mitochondrial dynamics towards pro-fusion state, and attenuate tumor growth in a mouse model of CRPC. In addition, to further investigate the molecular mechanism underlying the effects of exercise training on tumor growth, we employed a transcriptomics approach to identify intratumoral transcriptome and molecular pathways altered by voluntary wheel running.

## METHODS

### Cells and Reagents

We selected the aggressive CRPC cell line model, CWR-22RV1, known for its resistance to AR-targeted therapies such as enzalutamide (29, 30), to generate xenograft tumors in castrated mice. Human CWR-22RV1 (22RV1) cell line was obtained from the American Type Culture Collection (ATCC, Rockville, md, USA) and authenticated using short tandem repeat profiling, and tested for Mycoplasma contamination using the MycoAlert Mycoplasma Detection Kit (Lonza). Cells were cultured in RPMI-1640 medium supplemented with FBS (10%), penicillin (100 U/ml), streptomycin (100 μg/ml) and L-glutamine (300 μg/ml).

### Animals and Interventions

CRPC xenograft mouse model was established as previously described (31). In brief, male immunodeficient mice (SCID) were obtained from Taconic at 6-week of age. After one week of acclimatization, the mice were surgically castrated. Tumor cell inoculation occurred three days after castration for mice to recover from the surgery (Figure 1). Briefly, 22RV1 cells were resuspended in serum-free RPMI-1640 medium and mixed in a 1:1 ratio with Matrigel (BD Biosciences) prior to subcutaneous inoculation (1×10^6^ cells per injection) on flanks of both sides in castrated SCID mice. Twenty-four hours after tumor inoculation, mice were randomly assigned to either exercise training group using voluntary wheel running (VWR) or remained sedentary (SED) for 3 weeks (n=6/group) (Figure 1). Exercise training was performed by voluntary wheel running using a cage wheel running system with Hall Effect Sensors connected to a six-channel wheel counter interface and MDI Multi Interface Software (Columbus Instruments, Columbus, OH). Throughout the intervention, all mice were single housed. Tumor length (L) and width (W) were measured weekly by caliper, and tumor volumes were calculated (L x W^2^/2) and the average of the tumors from both sides was calculated. All mice were housed in a temperature-controlled (65–75°F) environment and maintained on a 12:12 hour light-dark cycle with food and water provided ad libitum. All animal procedures used on the mice were approved the Institutional Animal Care and Use Committee (IACUC) at the University of Massachusetts Boston (Protocol#: 2019156)

**FIGURE 1.**
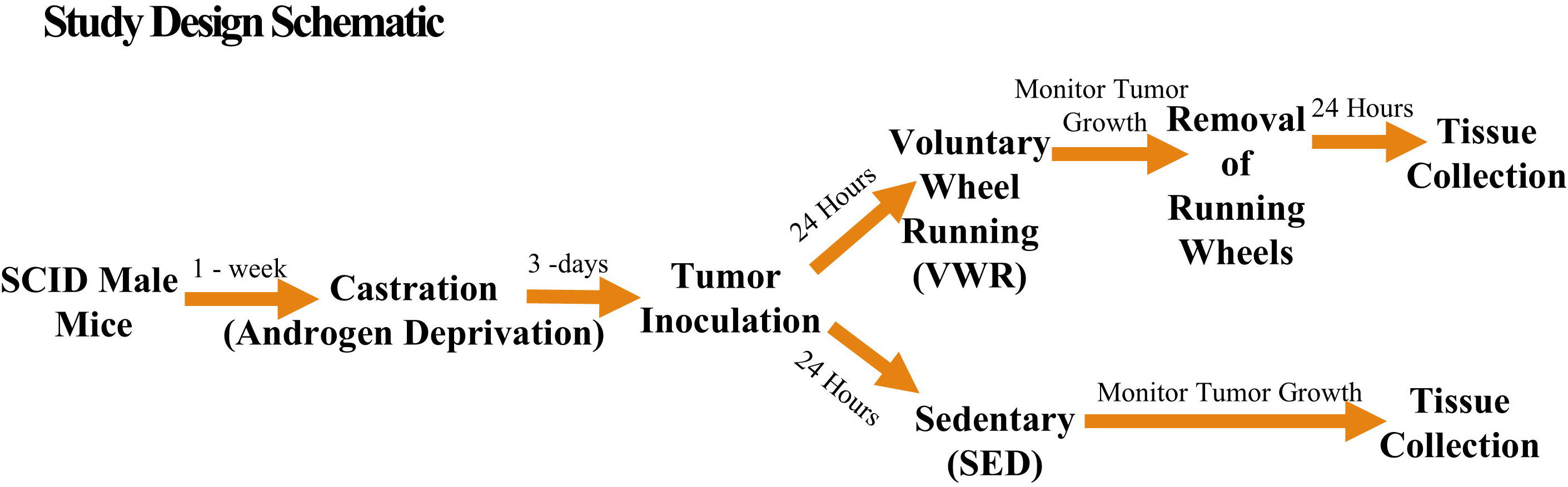
Schematic of Study Design.

### Tissue Collection

Twenty-four hours after the running wheel removal, animals were fasted overnight (12 h) before tissue collection. Euthanasia was performed using CO_2_ asphyxiation/cervical dislocation. Tumor tissues were collected, weighed, and snap-frozen in liquid nitrogen. Tissues were stored in –80°C until further analysis.

### Quantitative real-time PCR (RT-PCR)

RNA was extracted using RNeasy kit (Qiagen, Hilden, Germany) as previously described (32). Concentrations and purity of RNA samples were assessed on a NanoDrop One Microvolume UV-Vis Spectrophotometer (Thermo Fisher Scientific, Waltham, MA). cDNA was reversed transcribed from a 100ng of RNA using a High-Capacity cDNA Reverse Transcription Kits (Applied Biosystems, Foster City, CA) following manufacture instructions. cDNA was amplified in a 0.2 mL reaction containing TaqMan Universal PCR Master Mix, TaqMan Gene Expression Assay and RNase-free water. RT-PCR was performed using a QuantStudio 3 Real-Time PCR (Thermo Fisher Scientific, Waltham, MA), and results were analyzed using Design and Analysis Application (Thermo Fisher Scientific, Waltham, MA). Gene expression was quantified for all genes of interest (Supplementary Table 1) using the ΔΔCT method. The expression GAPDH was used as the housekeeping gene.

### Immunoblotting

Tumor tissues were homogenized as previously described (33). Protein concentrations from tumors were determined by a Pierce BCA Protein Assay Kit (Thermo Fisher Scientific, Waltham, MA), and equal amounts of protein were prepared to SDS-Page using a 4-20% gradient polyacrylamide gel (Bio-Rad, Hercules, CA) and transferred to a nitrocellulose membrane. After blocking, membranes were probed with a primary antibody and rocked overnight in 4° C. After washing with TBST membranes were then probed with an IRDye secondary antibody (Li-Cor, Lincoln, NE) and quantified using the Odyssey CLx software (Li-Cor, Lincoln, NE). All data were normalized to GAPDH protein expression. Immunoblot imaging was conducted on Licor. Supplemental Table 2 includes a complete list of all the suppliers, catalog numbers, and dilutions for primary and secondary antibodies used in this study.

### RNA Sequencing and Bioinformatics

RNA-seq library was prepared using TruSeq Stranded RNA LT Kit (Illumina). Sequencing was performed on NextSeq 2000 Illumina Genome Analyzer. The single-end reads were processed by FastQC and aligned by STAR (version 2.5.3a) to the human Ensemble genome (Ensembl, hg38/GRCh38) with all default configurations (34). Gene expression counts were generated using the subread-feature counts pipeline (version 1.6.2) (35, 36).

Gene expression analysis was then conducted using edgeR (version 3.24.3) (37), which uses Bayes estimation and exact tests based on the negative binomial distribution. Gene expression signals were logarithmically transformed (to base 2). Differentially expressed genes were determined based on the adjusted p-value threshold of 0.05 and log2 fold change threshold of abs (1.5) for up and down-regulated genes (Supplemental Table 3). The pre-ranked gene lists were used to conduct Gene Set Enrichment Analysis (GSEA) by using R package fgsea (version 1.22.0). The top pathways with normalized enrichment scores (NES) ranked by P value were plotted for visualization.

Gene ontology (GO) enrichment analysis was performed using g:Profiler (version e97_eg44_p13_d22abce) with g:SCS multiple testing correction method. A significance threshold of 0.05 was set (38).

### Statistical Analysis

All data are expressed as mean ± SEM. Student t-test was used to assess statistical differences between groups. Tumor growth from baseline was analyzed using One-way repeated measures of analysis of variance (ANOVA). All statistical analyses were performed using SPSS statistical software (27.0; SPSS, Inc, Chicago, IL). * P < 0.05; ** P < 0.01; *** P < 0.001.

## RESULTS

### Voluntary wheel running attenuates tumor growth in a mouse model of CRPC

There were no differences in body weight during the time course of the experiment, final body weight, muscle and heart weights between VWR and SED groups (Figure 2A and B, Supplemental Table 4). Epididymal fat pat weight was significantly lower in VWR group than SED group (P<0.05, Supplemental Table 4). Tumor progression was confirmed in mice from both VWR and SED groups (main effect of time, P < 0.001). Importantly, voluntary wheel running inhibited tumor formation as evidenced by smaller tumor volume at the initial time point (Day 10. Figure 2C, P = 0.048) in VWR group compared to SED group. In addition, three weeks of voluntary wheel running attenuated tumor progression throughout the time course (time x group interaction. P = 0.042, Figure 2C). Finally, there was a trend of significantly smaller tumor volume at the endpoint in VWR group compared to SED group (388.9±109.6 vs. 565.6±173.3 mm^3^, P = 0.061, Figure 2D). Consistently, the PCa diagnosis marker Prostate-Specific Antigen gene (also called *KLK3*) expression was significantly lower in tumor tissue collected from VWR group than SED group (1.3±0.3 vs. 1.8±0.1, P = 0.004, Figure 2E).

**FIGURE 2.**
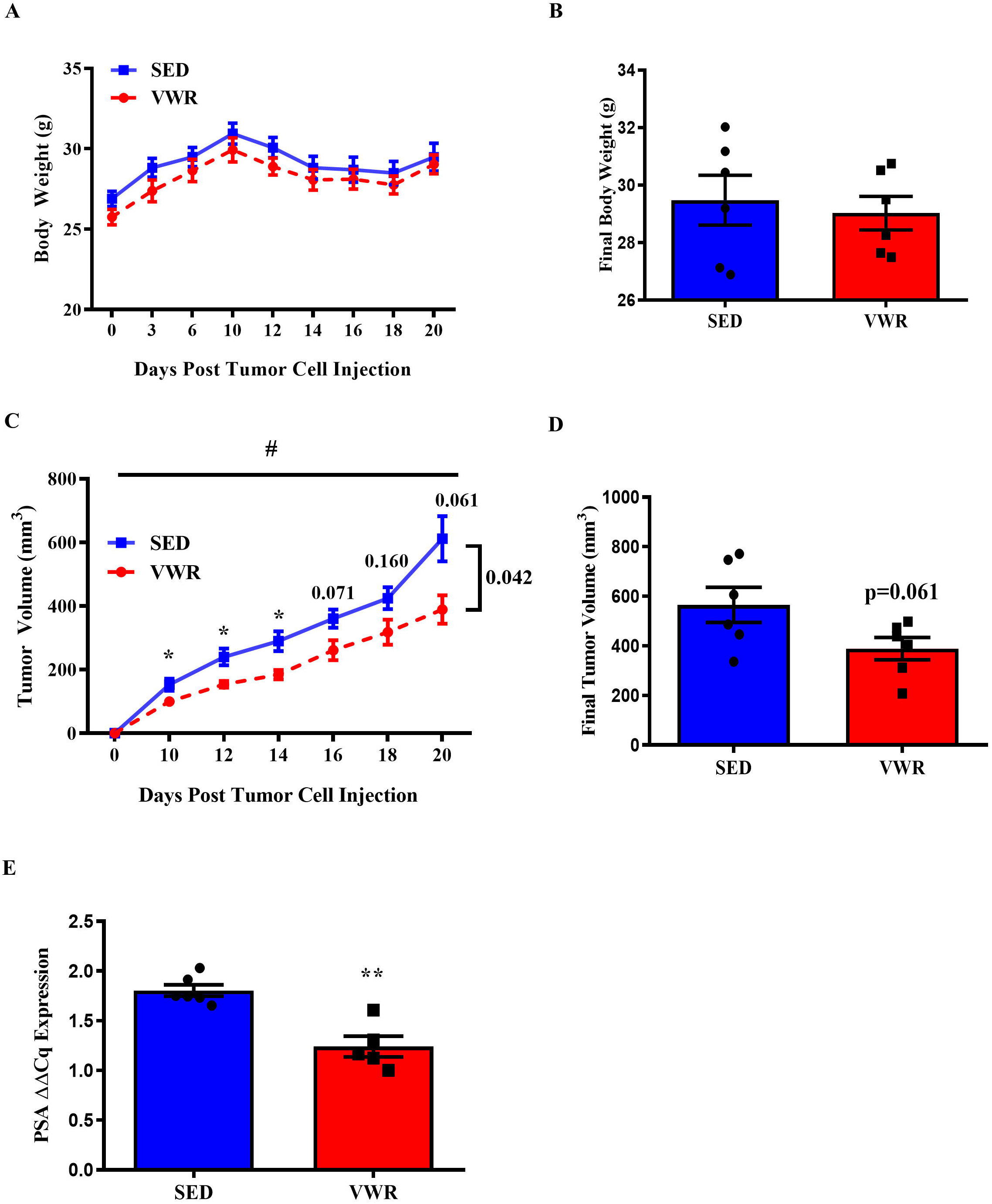
Voluntary wheel running attenuates tumor growth in a mouse model of CRPC. (A) Time course changes in body weight post tumor graft. (B) Final body weight. (C) Time course changes in tumor volume (mm^3^) post graft. (D) Final tumor volume (mm^3^). (E) PSA mRNA expression. N = 5-6/group. *P < 0.05; **P < 0.01 vs. SED. #P < 0.05 main effect of time, Data are expressed as Mean ± SEM.

### Voluntary wheel running modulates gene expression of DNA replication and AR signaling targets

The regulatory genes for cellular proliferation and DNA replication *MCM2*, *MCM6*, and *MCM7* were downregulated in the VWR group compared to SED (1.2±0.1 vs 2.1±0.1, P < 0.001; 1.4±0.1 vs. 2.2±0.2, P = 0.005; 1.1±0.1 vs. 2.1±0.2, P < 0.001; Figure 3A-C). Consistently, voluntary wheel running also resulted in downregulation of typic AR signaling target genes, *ELOVL5* and *FKBP5* (39), compared to the SED group (1.1±0.0 vs. 1.5±0.0, P < 0.001; 1.9±0.2 vs. 1.3 ±0.2, P = 0.001. Figure 3D and E).

**FIGURE 3.**
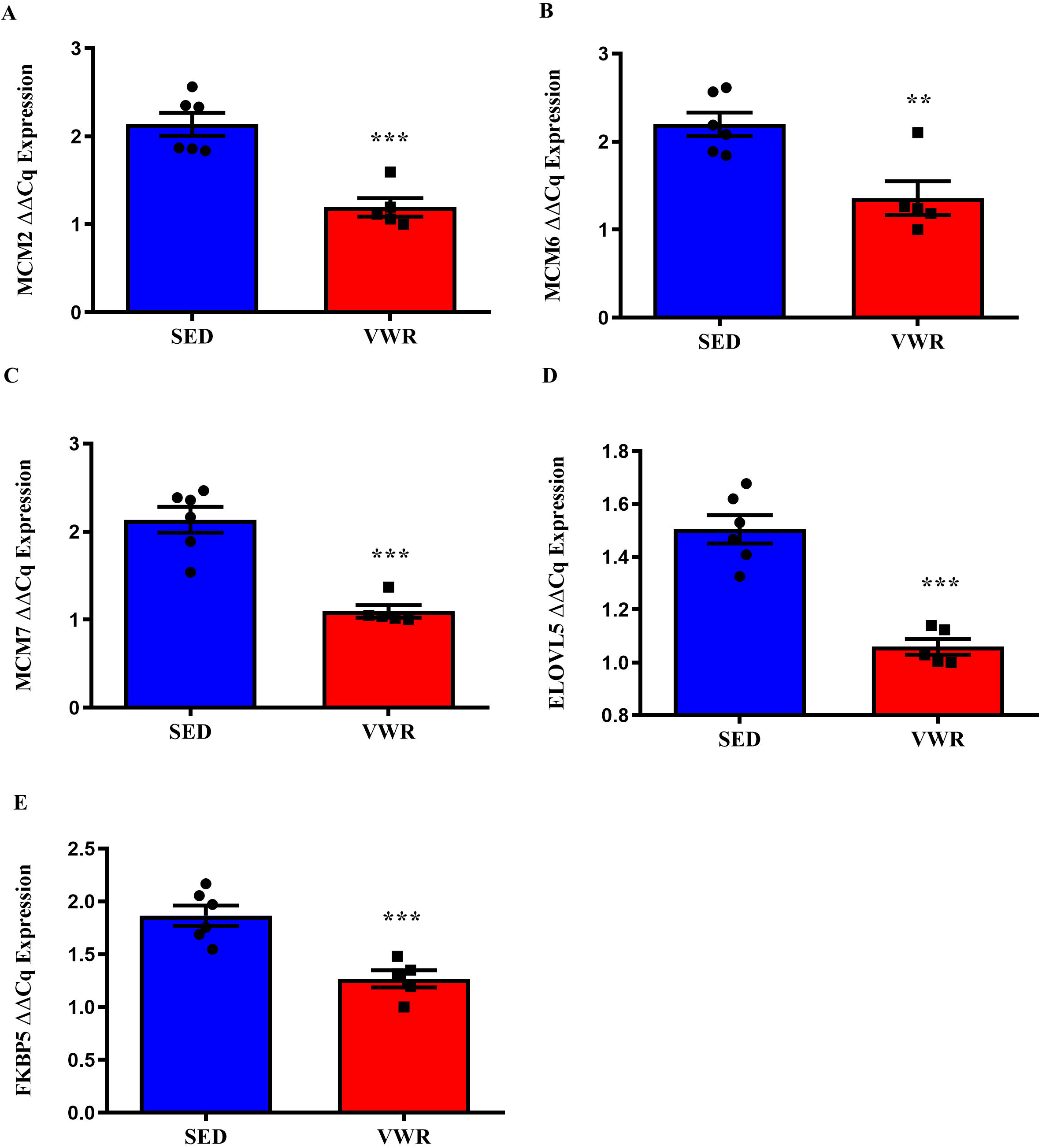
Voluntary wheel running modulates gene expression of cellular processes in tumor tissue from a mouse model of CRPC. (A) *MCM2* mRNA expression. (B) *MCM6* mRNA expression. (C) *MCM7* mRNA expression. (D) *ELOVL5* mRNA expression. (E) *FKBP5* mRNA expression. N = 5-6/group. **P < 0.01; ***P < 0.001 vs. SED. Data are expressed as Mean ± SEM.

### Voluntary wheel running alters transcription of mitochondrial dynamics regulatory markers

mRNA expression of *DNM1L* (The gene that encodes Drp1) and its associated adaptor *FIS1*, main regulator of mitochondrial fission process, were markedly lower in VWR group in comparison to SED group (1.5±0.2 vs. 2.8±0.5 and 1.4±0.2 vs. 2.5±0.4, P = 0.03 and 0.056, respectively. Figure 4A and B). In addition, mRNA expression of mitochondrial fusion markers *MFN1* and *OPA1* were also significantly reduced in mice who completed voluntary wheel running when compared to their sedentary counterparts (1.4±0.3 vs. 2.9±0.3 and 1.3±0.2 vs. 2.4±0.1, P = 0.01 and 0.001, respectively. Figure 4C and E). There was a trend of downregulation *in MFN2* mRNA expression in VWR group in comparison to SED group (1.5±0.2 vs. 2.1±0.2. P = 0.059. Figure 4D).

**FIGURE 4.**
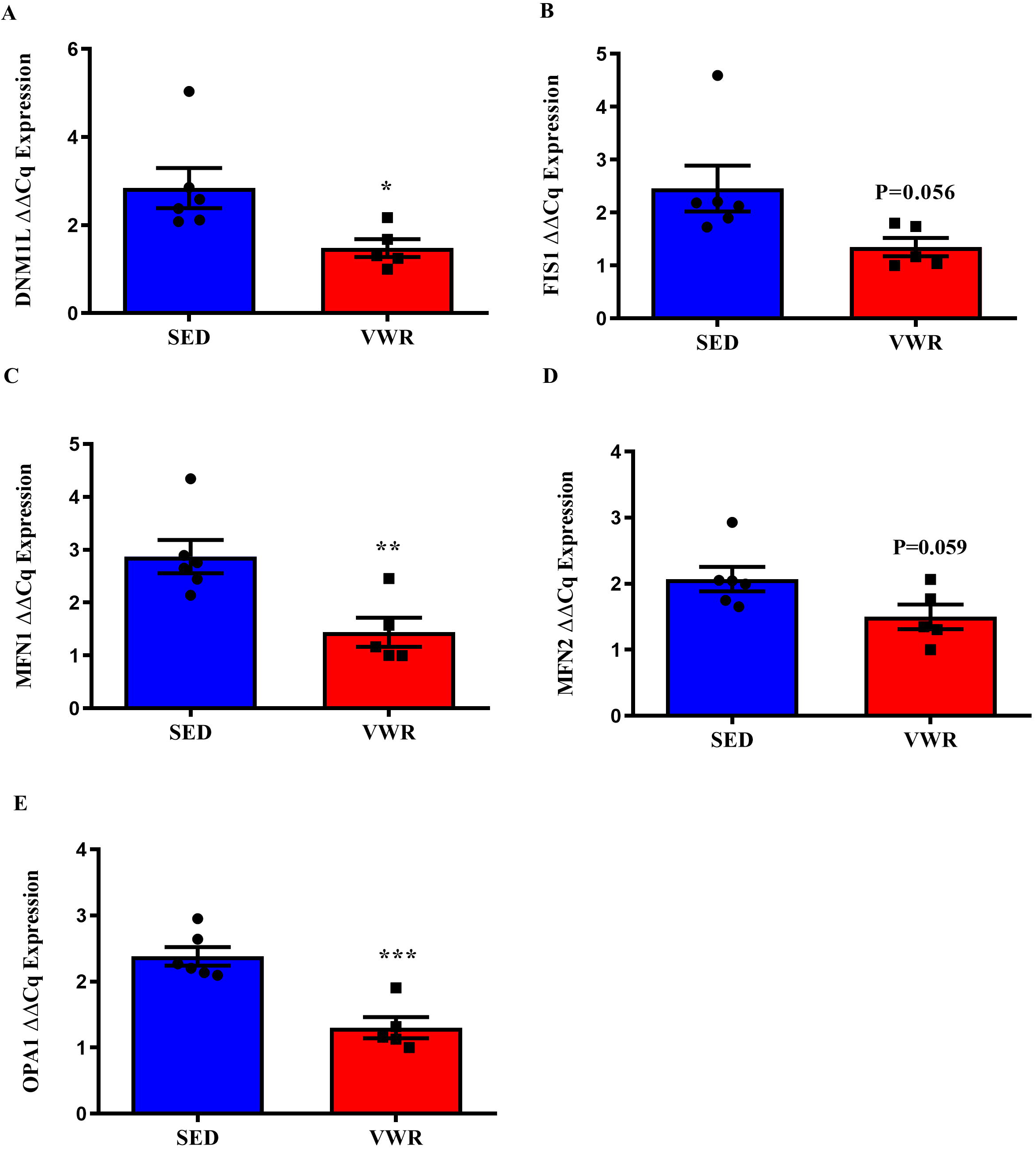
Voluntary wheel running alters gene expressions of regulatory markers of mitochondrial dynamics in tumor tissue from a mouse model of CRPC. (A) *DNM1L* mRNA expression. (B) *FIS1* mRNA expression. (C) *MFN1* mRNA expression. (D) *MFN2* mRNA expression. (E) *OPA1* mRNA expression. P < 0.05; **P < 0.01; ***P < 0.001 vs SED. Data are expressed as Mean ± SEM.

We next assessed protein expression of the regulatory markers for mitochondrial quality control and content. Surprisingly, there were no significant difference in protein expression of regulatory markers of mitochondrial dynamics (Drp1, FIS1, Mid51, MFN2, and OPA1), or autophagy (LC3B and P62) between the two groups (Figure 5A-G). There were also no statistical differences in mitochondrial content markers VDAC, citrate synthase, and OXPHOS between VWR and SED (Supplemental Figure 1A-D). Moreover, there was no difference in citrate synthase activity between the two groups (Supplemental Figure 1E).

**FIGURE 5.**
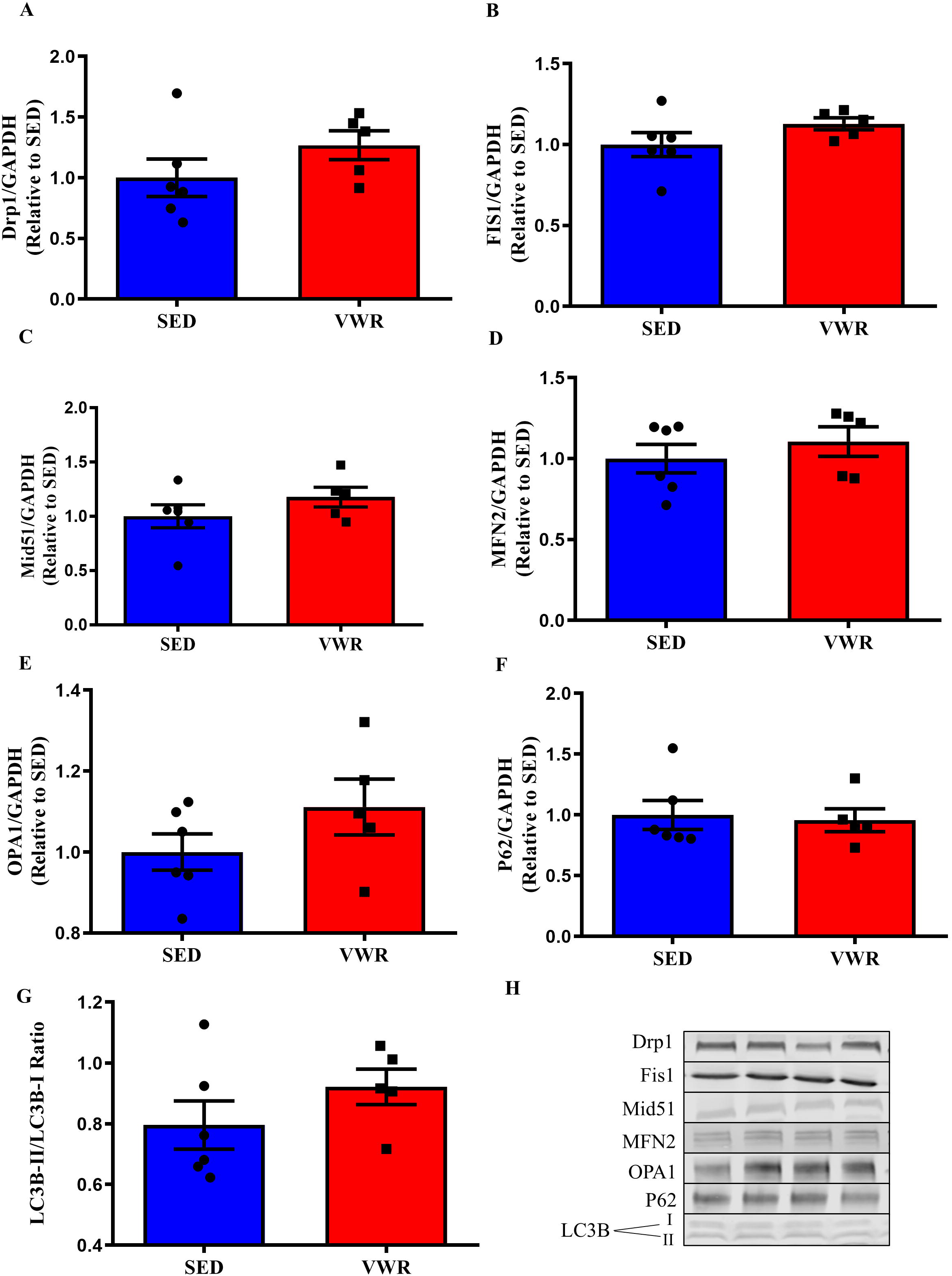
Voluntary wheel running did not change protein expression of regulatory markers of mitochondrial quality control in tumor tissue from a mouse model of CRPC. (A) Drp1 protein content. (B) Fis1 protein content. (C) Mid51 protein content. (D) MFN2 protein content. (E) OPA1 protein content. (F) P62 protein content. (G) LC3B II:I Ratio. (H) Representative immunoblots for proteins in A-G. N = 5-6/group. Data are expressed as Mean ± SEM.

### RNA sequencing data reveals global effects on gene expression and pathways in tumor tissue from voluntary wheel running mice

RNA sequencing revealed 287 differently expressed genes (157 up and 130 down) in tumor tissues collected from mice that completed three weeks of voluntary wheel running in comparison to their sedentary counterparts (Figure 6A and Supplemental Figure 3). The top 15 genes that were upregulated and downregulated in VWR group vs. SED group were presented in Table 1.

**FIGURE 6.**
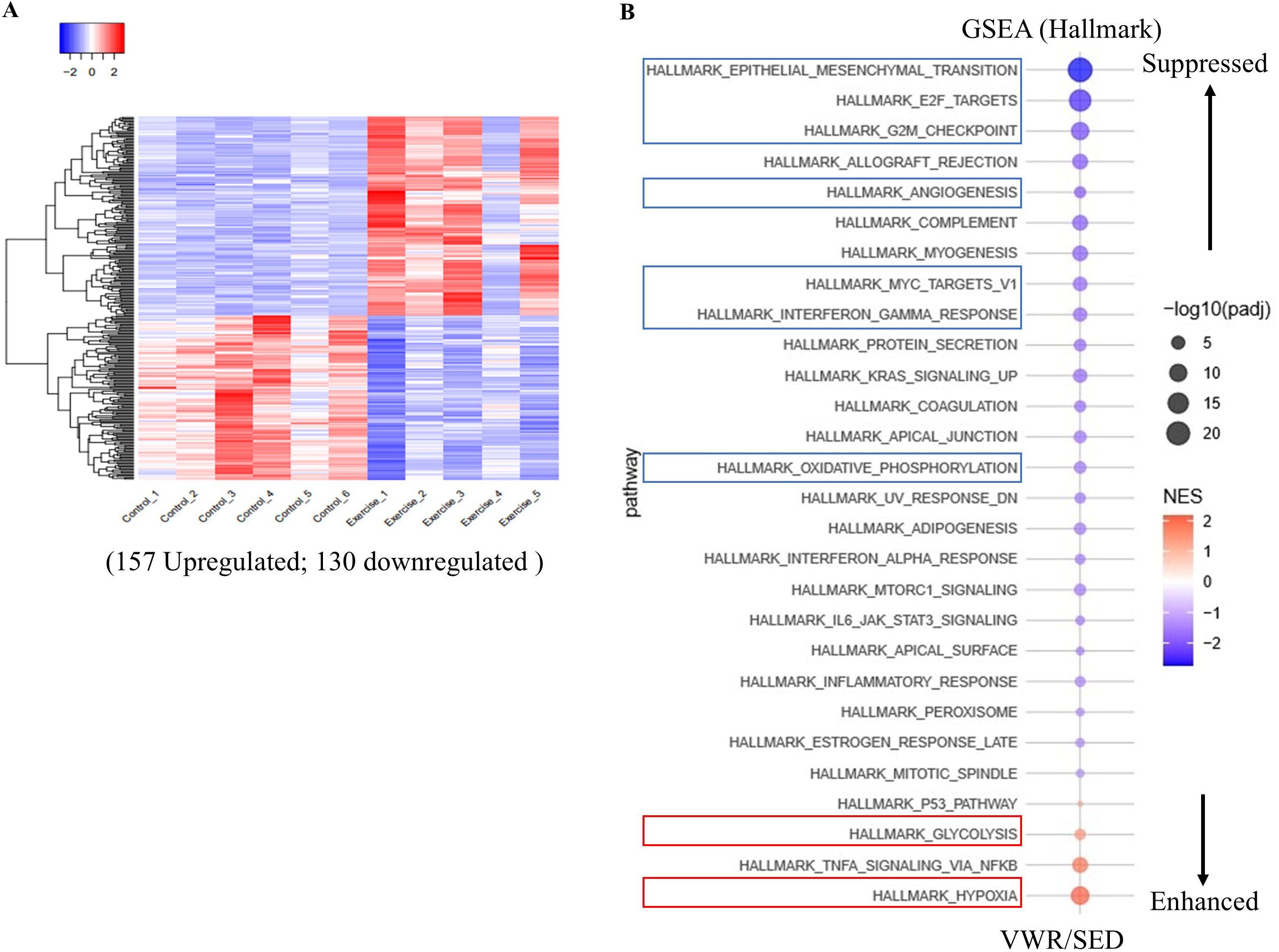
RNA sequencing data. (A) Heatmap view of differentially expressed genes between VWR and SED groups. (B) Gene Set Enrichment Analysis (GSEA) using Hallmark gene sets for the differentaily enriched pathways. Red indicates higher expression and blue indicates lower expression. N=5-6/group.

**Table 1.** Top 15 genes upregulated and downregulated by voluntary wheel running.

| **Gene ID** | **Gene Name** | **Fold Change** | **P-adj** |
| --- | --- | --- | --- |
| **Upregulated** |  |  |  |
| **SPAG4** | Sperm associated antigen 4 | 1.88 | 0.001 |
| **PER1** | Period circadian regulator 1 | 1.86 | 0.001 |
| **FOS** | Fos proto-oncogene, AP-1 transcription factor subunit | 1.85 | 0.001 |
| **COL7A1** | Collagen type VII alpha 1 chain | 1.81 | 0.001 |
| **ANKRD37** | Ankyrin repeat domain 37 | 1.77 | 0.001 |
| **MAFF** | MAF bZIP transcription factor F | 1.76 | 0.001 |
| **CCDC183** | Coiled-coil domain containing 183 | 1.76 | 0.001 |
| **LOC646471** | Uncharacterized LOC646471 | 1.75 | 0.001 |
| **SH3D21** | SH3 domain containing 21 | 1.75 | 0.001 |
| **HDAC10** | Histone deacetylase 10 | 1.75 | 0.001 |
| **CAPN5** | Calpain 5 | 1.74 | 0.001 |
| **DPYSL4** | Dihydropyrimidinase like 4 | 1.74 | 0.001 |
| **PPFIA4** | PTPRF interacting protein alpha 4 | 1.73 | 0.001 |
| **KRTAP5-1** | Keratin associated protein 5-1 | 1.73 | 0.001 |
| **IDUA** | Alpha-L-iduronidase | 1.72 | 0.001 |
| **Downregulated** |  |  |  |
| **VCAM1** | Vascular cell adhesion molecule 1 | -2.03 | 0.001 |
| **EPB41L3** | Erythrocyte membrane protein band 4.1 like 3 | -1.98 | 0.001 |
| **SEMA3A** | Semaphorin 3A | -1.89 | 0.001 |
| **SRPX** | Sushi repeat containing protein X-linked | -1.87 | 0.001 |
| **PABPC3** | Poly(A) binding protein cytoplasmic 3 | -1.86 | 0.001 |
| **COL8A2** | Collagen type VIII alpha 2 chain | -1.83 | 0.001 |
| **ZNF521** | Zinc finger protein 521 | -1.83 | 0.001 |
| **EDIL3** | EGF like repeats and discoidin domains | -1.82 | 0.001 |
| **LUM** | Lumican | -1.81 | 0.001 |
| **CACNA1G** | Calcium voltage-gated channel subunit alpha1 G | -1.81 | 0.001 |
| **EBF3** | EBF transcription factor 3 | -1.79 | 0.001 |
| **APBB1IP** | Amyloid beta precursor protein binding family B member 1 interacting protein | -1.78 | 0.001 |
| **GJA4** | Gap junction protein alpha 4 | -1.78 | 0.001 |
| **ETS1** | ETS proto-oncogene 1, transcription factor | -1.77 | 0.001 |
| **PDIA3P1** | Protein disulfide isomerase family A member 3 pseudogene 1 | -1.77 | 0.001 |

Gene Set Enrichment Analysis (GSEA) using Hallmark gene sets on the RNA-seq data revealed that several pathways commonly associated with CRPC tumor growth were downregulated in tumors collected from VWR mice compared to the SED group (Figure 6B). These pathways include cell cycle regulation (E2F targets, G2M checkpoint and MYC targets), epithelial-mesenchymal transition, angiogenesis and oxidative phosphorylation. In contrast, only a few pathways, such as glycolysis and hypoxia, were downregulated (Figure 6B). Consistent with GSEA analysis, gene ontology pathway analysis revealed downregulation of some top pathways related to angiogenesis in tumors from VWR mice (angiogenesis, extracellular matrix organization and endothelial cell proliferation) and upregulation of several pathways, including glycolytic process, Histone H3-K9 demethylation, response to hypoxia and apoptotic process (Table 2).

**Table 2.**
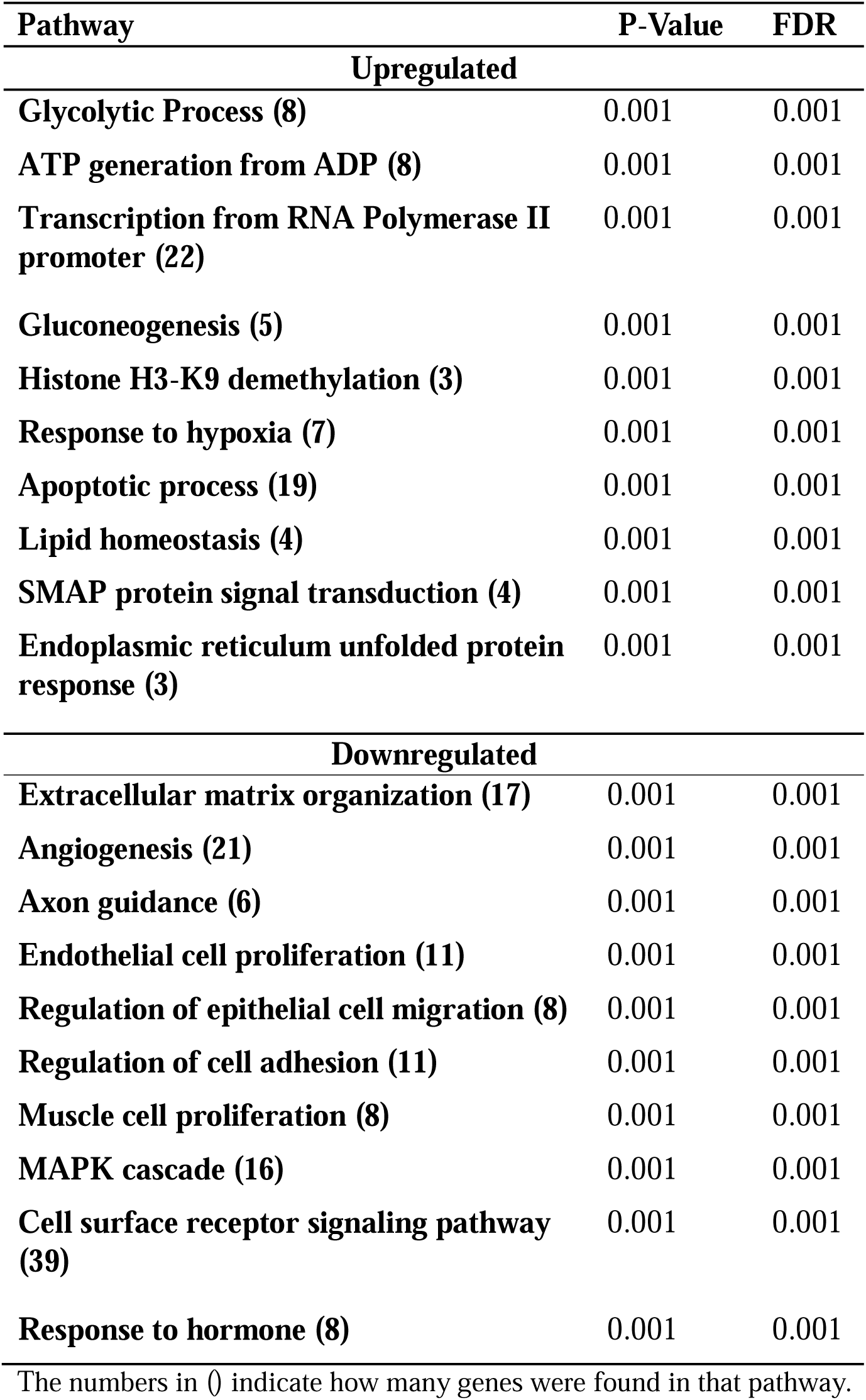
Top 10 pathways altered by voluntary wheel running.

| **Pathway** | **P-Value** | **FDR** |
| --- | --- | --- |
| **Upregulated** |  |  |
| **Glycolytic Process (8)** | 0.001 | 0.001 |
| **ATP generation from ADP (8)** | 0.001 | 0.001 |
| **Transcription from RNA Polymerase II promoter (22)** | 0.001 | 0.001 |
| **Gluconeogenesis (5)** | 0.001 | 0.001 |
| **Histone H3-K9 demethylation (3)** | 0.001 | 0.001 |
| **Response to hypoxia (7)** | 0.001 | 0.001 |
| **Apoptotic process (19)** | 0.001 | 0.001 |
| **Lipid homeostasis (4)** | 0.001 | 0.001 |
| **SMAP protein signal transduction (4)** | 0.001 | 0.001 |
| **Endoplasmic reticulum unfolded protein response (3)** | 0.001 | 0.001 |
| **Downregulated** |  |  |
| **Extracellular matrix organization (17)** | 0.001 | 0.001 |
| **Angiogenesis (21)** | 0.001 | 0.001 |
| **Axon guidance (6)** | 0.001 | 0.001 |
| **Endothelial cell proliferation (11)** | 0.001 | 0.001 |
| **Regulation of epithelial cell migration (8)** | 0.001 | 0.001 |
| **Regulation of cell adhesion (11)** | 0.001 | 0.001 |
| **Muscle cell proliferation (8)** | 0.001 | 0.001 |
| **MAPK cascade (16)** | 0.001 | 0.001 |
| **Cell surface receptor signaling pathway (39)** | 0.001 | 0.001 |
| **Response to hormone (8)** | 0.001 | 0.001 |

The numbers in () indicate how many genes were found in that pathway.

## DISCUSSION

It is crucial to improve the quality of life while control CRPC tumor progression. In this study, we examined the impact of voluntary exercise training on tumor growth and the regulatory machinery of mitochondrial dynamics in a preclinical model of CRPC implanted with human prostate cancer cells. The main finding of the present study was that voluntary wheel running was effective in delaying tumor initiation and growth, which coincided with reduced mRNA expressions of regulatory proteins of mitochondrial dynamics in mice with CRPC. We further sought to identify molecular regulators and pathways that may explain the beneficial effects of exercise training on tumor progression in mice with CRPC. Strikingly, our RNA-seq analysis using tumor tissues from CRPC xenograft tumors revealed that three weeks of voluntary wheel running resulted in significant downregulation of multiple molecular pathways/processes related to cell proliferation (E2F Targets and MYC targets), and angiogenesis (Extracellular matrix organization, Angiogenesis, and Endothelial cell proliferation). Together, these data highlight the effectiveness of exercise training on controlling tumor progression in CRPC and shed light on potential molecular mechanisms underlying these beneficial effects.

One of the major findings of the present study is that voluntary exercise training inhibited the initiation and progression of PCa tumors in a preclinical model of CRPC implanted with human prostate cancer cells. 22RV1 cells are a human prostate cancer cell line widely used as a standard CRPC model due to their high expression of the androgen receptor splice variant 7 (AR-V7) and are commonly used to test potential therapeutic strategies for treating CRPC (40). It has been shown that 22RV1 cells are resistant to abiraterone acetate and enzalutamide, key AR-targeted therapeutic agents for treating CRPC (29). In line with this, our study found that voluntary exercise training effectively reduced expression of AR/AR-V7 downstream pathway targets (i.e., *ELOVL5* and *FKBP5*), which may be responsible for the attenuated tumor progression in our model of CRPC. Together, our findings of reduced tumor growth and AR-V7 signaling by voluntary exercise training in a xenograft model of CRPC using 22RV1 cells is of critical clinical significance as it provides evidence for developing exercise regimens to treat CRPC. Our findings are consistent with a large body of literature demonstrating that voluntary exercise training is effective in delaying tumor onset and progression in rodent models of PCa (41–45). In addition, our finding was partly in agreement with a recent study using the same mouse model of CRPC (11). Recently, Yang et al., reported that voluntary wheel running inhibited tumor progression in a mouse cell line model of CRPC, but failed to observe any difference in tumor initiation (11). The discrepancy in the effects of voluntary wheel running on tumor initiation may be due to the differences in cell lines used for tumor implantation (mouse (RM1) vs. human PCa cell line (22RV1)), implanted tumor cell volume (2×10^6^ vs. 1×10^6^ cells), and/or total time exposed to the running wheel (1 hour vs. 24 hours). Overall, our findings corroborated with previous studies demonstrating the benefits of exercise on attenuating PCa tumor growth and progression (41–45) but further extend these findings by demonstrating such benefits in an advanced PCa mouse model (i.e., CRPC) using a human prostate carcinoma cell line, a more clinical relevant CRPC model.

In the present study, we observed that voluntary wheel running downregulates mRNA expression of Minichromosome maintenance proteins (MCMs). These MCMs are essential for DNA replication for maintaining normal cellular processes, especially during the DNA initiation and elongation stages (46, 47). Due to their vital role in genome duplication in proliferating cells, overexpression of MCMs are commonly observed in tumor formation and growth (48). The lower mRNA expressions of MCMs in tumors collected from exercise trained mice suggests that voluntary wheel running may delay tumor growth by inhibiting tumor cell proliferation.

Although we did not directly measure cell proliferation, the GSEA and GO pathway analyses from our RNA sequencing data identified significant downregulation of the cell cycle regulation pathways, such as E2F targets and MYC targets. In addition, findings from a recent study indeed reported that voluntary wheel running inhibited cell proliferation in a mouse model of CRPC (11), further corroborating our findings on the effects of exercise on tumor cell proliferation.

Accumulating evidence supports the notion that mitochondrial metabolism (i.e., oxidative phosphorylation) is elevated in CRPC tumors, which plays an important role in tumorigenesis and tumor progression (13–15). The orchestrated balance between mitochondrial fission and fusion is crucial for maintaining mitochondrial morphology and function (19). Dysregulated mitochondrial dynamics has been linked with cancer progression and metastasis (21, 22). In the present study, among changes in all regulatory machinery of mitochondrial dynamics, reduced expression of mitochondrial fission mediator, Drp1, in tumors collected from VWR mice is of most interest. Genomics analyses from clinical datasets have shown that Drp1 mRNA expression was consistently higher in CRPC tumors from patients than normal prostate tissues (49–51). In addition, Lee et al. recently reported that androgen signaling promoted mitochondrial metabolism by upregulating Drp1 expression and inhibiting Drp1 induced a metabolic stress response resulting in attenuation of androgen receptor-mediated growth of PCa (15). These findings are consistent with our study, suggesting that voluntary exercise training may inhibit mitochondrial metabolism via reducing Drp1-medaited mitochondrial fission. Although PCa tumor cells, like other cancer cells, preferentially rely on glycolysis to produce ATP, emerging evidence has shown that PCa cells that are resistant to enzalutamide treatment (e.g., 22RV1) switch from glycolysis to oxidative phosphorylation and have increased mitochondrial metabolism (e.g., TCA cycle and oxidative phosphorylation) to meet the energy needs for the uncontrolled cell proliferation and metastasis (13, 52). More important, Basu et al., reported that treating 22RV1 PCa cells with mitochondrial metabolic inhibitors effectively inhibited PCa cell proliferation (52). While we did not measure mitochondrial metabolism, our results of reduced expression of both mitochondrial fission (e.g., Drp1 and Fis1) and fusion (MFN1, MFN2 and OPA1) regulators, coupled with downregulated oxidative phosphorylation pathway from GSEA analysis, suggest that mitochondria were less active in tumors collected from VWR mice, which supports the hypothesis that voluntary exercise training may reduce mitochondrial metabolism. However, whether the reduced mitochondrial activity/metabolism was the cause or consequence of reduced tumor cell proliferation following voluntary exercise training remains unknown.

Contrary to the data at the transcription level, we did not find consistent changes of these regulatory markers of mitochondrial dynamics at the protein level. The inconsistency between mRNA and protein expressions of markers of mitochondrial dynamics may be due to the dynamic nature of mitochondrial dynamics and the “snapshot” approach may have missed the window of the alterations in mitochondrial dynamics regulatory markers at the protein level. This is corroborated by previous studies where differential patterns of mRNA and protein expressions were reported following voluntary wheel running (53). Future studies should further investigate the posttranslational modifications of these regulatory markers of mitochondrial dynamics (e.g., Drp1), as well as morphological changes in tumor tissues to provide definitive evidence on the effect of exercise training on mitochondrial dynamics alterations in tumor tissues.

We further conducted RNA sequencing on tumor tissues obtained from SED and VWR groups to identify key genes and pathways that may underlie the aforementioned beneficial effects of exercise training on CRPC tumor. The top-ranking pathways include several that are associated with angiogenesis, such as downregulation of extracellular matrix organization, angiogenesis and endothelial cell proliferation. Angiogenesis is a fundamental process in PCa tumor progression and has been recommended as a therapeutic target for treating CRPC (54). Angiogenesis has been reported to promote endothelial cell proliferation, which in turn, increases resistance to apoptosis. Our findings of downregulated endothelial cell proliferation pathway and upregulation of apoptosis pathway suggest that VWR may have delayed tumor progression by downregulating angiogenesis and subsequently inducing apoptosis of tumor cells. In agreement, Zielinski, et. al., reported that exercise reduced tumor blood vessel density (55). In contrast, several studies found that exercise training enhanced angiogenesis in rodent models of PCa, including improvement in microvasculature and enhanced blood flow (56, 57). The disagreement between our study and these studies may be due to the progression of tumor, heterogeneity of tumor microenvironment, and the differences with murine models. One limitation of our study is that we did not measure angiogenesis or vascularization. Future studies should be warranted to evaluate vasculature in tumor samples collected from CRPC mice following VWR to validate our findings.

Interestingly, in the present study, glycolysis and hypoxia pathways were elevated while oxidative phosphorylation was downregulated in tumor tissues collected from VWR mice in comparison to SED mice. This is unexpected due to the fact that glycolysis typically promotes tumor progression (58) and is elevated in patients with CRPC compared with those having primary PCa tumors and normal controls (59). We speculate that the elevated pathways in glycolysis and hypoxia may be a secondary adaptation in response to the decreased angiogenesis and mitochondrial activity in tumor microenvironment. Over time, in order to meet the energy demand, CRPC tumor cells may have adapted to the environment by increasing glycolysis and hypoxia pathway. As mentioned above, some cancer cells, including PCa cells that are resistant to enzalutamide treatment, switch from glycolysis to oxidative phosphorylation during invasive state to meet their high energy demand (52, 60, 61). Therefore, such metabolic reprogramming in tumor tissue following voluntary exercise training may be an unfavorable adaptation for CPRC tumor cells that are forced to rely on glycolysis with insufficient energy generation, which may eventually lead to reduced cell proliferation and tumor growth. Future studies are warranted to investigate metabolic functions, particularly mitochondrial metabolism to see whether glycolysis, oxidative phosphorylation, and TCA cycle are altered in CRPC tumors following exercise training.

Our study has limitations. First, although xenograft models are considered a representative cancerous model, the tumor microenvironment may not be as applicable as orthopedic models. Second, the exercise variables including intensity, duration, and volume were uncontrolled. Due to voluntary wheel running being a non-prescribed modality of exercise, these exercise parameters are not controlled. However, voluntary wheel running, in contrast to forced exercise methods such as treadmill running, is less stressful and may better reflect the natural exercise behavior of mice with CRPC, a more advanced form of the disease.

In conclusion, our study demonstrated that in a CRPC mouse model, three weeks of voluntary wheel running was effective in delaying tumor initiation and progression, which coincided with reduced transcription of regulatory markers of mitochondrial dynamics. We further identified downregulation of multiple molecular pathways/processes related to cell proliferation (E2F Targets and MYC targets) and angiogenesis (extracellular matrix organization, angiogenesis, and endothelial cell proliferation) that may be responsible for the delayed tumor initiation and progression following voluntary exercise training. Together, our study provides valuable information demonstrating that voluntary exercise training is an effective approach for reducing tumor progression in CRPC, a more aggressive stage of PCa and novel insights into the mechanisms by which exercise modulates CRPC recuring and progression.

## ACKNOWLEDGMENTS

This study was supported by the University of Massachusetts Boston (Startup Funds to K.Z.) and the National Institutes of Health (R15DK131512 to K.Z.; R01CA211350 to C.C.)

## REFERENCES

1. Rawla P. Epidemiology of Prostate Cancer. World J Oncol. 2019;10(2):63–89.

2. Harris WP, Mostaghel EA, Nelson PS, Montgomery B. Androgen deprivation therapy: progress in understanding mechanisms of resistance and optimizing androgen depletion. Nat Clin Pract Urol. 2009;6(2):76–85.

3. Philip A. Watson VKA, Charles L. Sawyers. Emerging Mechanisms of Resistance to Androgen Receptor Inhibitors in Prostate Cancer. Nat Rev Cancer. 2015:701–11.

4. Chung DY, Kang DH, Kim JW et al. Comparison of Oncologic Outcomes Between Two Alternative Sequences with Abiraterone Acetate and Enzalutamide in Patients with Metastatic Castration-Resistant Prostate Cancer: A Systematic Review and Meta-Analysis. Cancers (Basel). 2019;12(1).

5. Richman EL, Kenfield SA, Stampfer MJ, Paciorek A, Carroll PR, Chan JM. Physical activity after diagnosis and risk of prostate cancer progression: data from the cancer of the prostate strategic urologic research endeavor. Cancer Res. 2011;71(11):3889–95.

6. Galvao DA, Taaffe DR, Spry N et al. Enhancing active surveillance of prostate cancer: the potential of exercise medicine. Nat Rev Urol. 2016;13(5):258–65.

7. Keilani M, Hasenoehrl T, Baumann L et al. Effects of resistance exercise in prostate cancer patients: a meta-analysis. Support Care Cancer. 2017;25(9):2953–68.

8. Pak S, Kim MS, Park EY, Kim SH, Lee KH, Joung JY. Association of Body Composition With Survival and Treatment Efficacy in Castration-Resistant Prostate Cancer. Front Oncol. 2020;10:558.

9. Giovannucci EL, Liu Y, Leitzmann MF, Stampfer MJ, Willett WC. A prospective study of physical activity and incident and fatal prostate cancer. Arch Intern Med. 2005;165(9):1005–10.

10. Hart NH, Galvao DA, Newton RU. Exercise medicine for advanced prostate cancer. Curr Opin Support Palliat Care. 2017;11(3):247–57.

11. Yang Z, Gao Y, He K et al. Voluntarily wheel running inhibits the growth of CRPC xenograft by inhibiting HMGB1 in mice. Exp Gerontol. 2023;174:112118.

12. Vikramdeo KS, Sharma A, Anand S et al. Mitochondrial Alterations in Prostate Cancer: Roles in Pathobiology and Racial Disparities. Int J Mol Sci. 2023;24(5).

13. Fontana F, Anselmi M, Limonta P. Unraveling the Peculiar Features of Mitochondrial Metabolism and Dynamics in Prostate Cancer. Cancers (Basel). 2023;15(4).

14. Massie CE, Lynch A, Ramos-Montoya A et al. The androgen receptor fuels prostate cancer by regulating central metabolism and biosynthesis. EMBO J. 2011;30(13):2719–33.

15. Lee YG, Nam Y, Shin KJ et al. Androgen-induced expression of DRP1 regulates mitochondrial metabolic reprogramming in prostate cancer. Cancer Lett. 2020;471:72–87.

16. Held NM, Houtkooper RH. Mitochondrial quality control pathways as determinants of metabolic health. Bioessays. 2015;37(8):867–76.

17. Twig G, Elorza A, Molina AJ et al. Fission and selective fusion govern mitochondrial segregation and elimination by autophagy. EMBO J. 2008;27(2):433–46.

18. Loson OC, Song Z, Chen H, Chan DC. Fis1, Mff, MiD49, and MiD51 mediate Drp1 recruitment in mitochondrial fission. Mol Biol Cell. 2013;24(5):659-67.

19. Williams M, Caino MC. Mitochondrial Dynamics in Type 2 Diabetes and Cancer. Front Endocrinol (Lausanne). 2018;9:211.

20. Sedlackova L, Korolchuk VI. Mitochondrial quality control as a key determinant of cell survival. Biochim Biophys Acta Mol Cell Res. 2019;1866(4):575–87.

21. Ghosh P, Vidal C, Dey S, Zhang L. Mitochondria Targeting as an Effective Strategy for Cancer Therapy. Int J Mol Sci. 2020;21(9).

22. Ma Y, Wang L, Jia R. The role of mitochondrial dynamics in human cancers. Am J Cancer Res. 2020;10(5):1278–93.

23. Seo JH, Agarwal E, Chae YC et al. Mitochondrial fission factor is a novel Myc-dependent regulator of mitochondrial permeability in cancer. EBioMedicine. 2019;48:353–63.

24. Philley JV, Kannan A, Qin W et al. Complex-I Alteration and Enhanced Mitochondrial Fusion Are Associated With Prostate Cancer Progression. J Cell Physiol. 2016;231(6):1364–74.

25. Fealy CE, Mulya A, Lai N, Kirwan JP. Exercise training decreases activation of the mitochondrial fission protein dynamin-related protein-1 in insulin-resistant human skeletal muscle. J Appl Physiol (1985). 2014;117(3):239-45.

26. Yan Z, Lira VA, Greene NP. Exercise training-induced regulation of mitochondrial quality. Exerc Sport Sci Rev. 2012;40(3):159–64.

27. Trewin AJ, Berry BJ, Wojtovich AP. Exercise and Mitochondrial Dynamics: Keeping in Shape with ROS and AMPK. Antioxidants (Basel). 2018;7(1).

28. Stroh AM, Stanford KI. Exercise-induced regulation of adipose tissue. Curr Opin Genet Dev. 2023;81:102058.

29. Li Y, Chan SC, Brand LJ, Hwang TH, Silverstein KA, Dehm SM. Androgen receptor splice variants mediate enzalutamide resistance in castration-resistant prostate cancer cell lines. Cancer Res. 2013;73(2):483–9.

30. Bennett L, Jaiswal PK, Harkless RV et al. Glucocorticoid Receptor (GR) Activation Is Associated with Increased cAMP/PKA Signaling in Castration-Resistant Prostate Cancer. Mol Cancer Ther. 2024;23(4):552–63.

31. Li M, Liu M, Han W et al. LSD1 Inhibition Disrupts Super-Enhancer-Driven Oncogenic Transcriptional Programs in Castration-Resistant Prostate Cancer. Cancer Res. 2023;83(10):1684–98.

32. Kugler BA, Gundersen AE, Li J et al. Roux-en-Y gastric bypass surgery restores insulin-mediated glucose partitioning and mitochondrial dynamics in primary myotubes from severely obese humans. Int J Obes (Lond). 2020;44(3):684–96.

33. Kugler BA, Deng W, Duguay AL et al. Pharmacological inhibition of dynamin-related protein 1 attenuates skeletal muscle insulin resistance in obesity. Physiol Rep. 2021;9(7):e14808.

34. Dobin A, Davis CA, Schlesinger F et al. STAR: ultrafast universal RNA-seq aligner. Bioinformatics. 2013;29(1):15–21.

35. Liao Y, Smyth GK, Shi W. The Subread aligner: fast, accurate and scalable read mapping by seed-and-vote. Nucleic Acids Res. 2013;41(10):e108.

36. Liao Y, Smyth GK, Shi W. featureCounts: an efficient general purpose program for assigning sequence reads to genomic features. Bioinformatics. 2014;30(7):923–30.

37. Robinson MD, McCarthy DJ, Smyth GK. edgeR: a Bioconductor package for differential expression analysis of digital gene expression data. Bioinformatics. 2010;26(1):139–40.

38. Raudvere U, Kolberg L, Kuzmin I et al. g:Profiler: a web server for functional enrichment analysis and conversions of gene lists (2019 update). Nucleic Acids Res. 2019;47(W1):W191–W8.

39. Han W, Gao S, Barrett D et al. Reactivation of androgen receptor-regulated lipid biosynthesis drives the progression of castration-resistant prostate cancer. Oncogene. 2018;37(6):710–21.

40. Sramkoski RM, Pretlow TG, 2nd, Giaconia JM et al. A new human prostate carcinoma cell line, 22Rv1. In Vitro Cell Dev Biol Anim. 1999;35(7):403-9.

41. Esser KA, Harpole CE, Prins GS, Diamond AM. Physical activity reduces prostate carcinogenesis in a transgenic model. Prostate. 2009;69(13):1372–7.

42. Jones LW, Antonelli J, Masko EM et al. Exercise modulation of the host-tumor interaction in an orthotopic model of murine prostate cancer. J Appl Physiol (1985). 2012;113(2):263-72.

43. Patel DI, Abuchowski K, Bedolla R et al. Nexrutine and exercise similarly prevent high grade prostate tumors in transgenic mouse model. PLoS One. 2019;14(12):e0226187.

44. Zheng X, Cui XX, Gao Z et al. Inhibitory effect of dietary atorvastatin and celecoxib together with voluntary running wheel exercise on the progression of androgen-dependent LNCaP prostate tumors to androgen independence. Exp Ther Med. 2011;2(2):221–8.

45. Zheng X, Cui XX, Huang MT et al. Inhibitory effect of voluntary running wheel exercise on the growth of human pancreatic Panc-1 and prostate PC-3 xenograft tumors in immunodeficient mice. Oncol Rep. 2008;19(6):1583–8.

46. Tye BK. MCM proteins in DNA replication. Annu Rev Biochem. 1999;68:649–86.

47. Forsburg SL. Eukaryotic MCM proteins: beyond replication initiation. Microbiol Mol Biol Rev. 2004;68(1):109–31.

48. Lei M. The MCM complex: its role in DNA replication and implications for cancer therapy. Curr Cancer Drug Targets. 2005;5(5):365–80.

49. Varambally S, Yu J, Laxman B et al. Integrative genomic and proteomic analysis of prostate cancer reveals signatures of metastatic progression. Cancer Cell. 2005;8(5):393–406.

50. Yu YP, Landsittel D, Jing L et al. Gene expression alterations in prostate cancer predicting tumor aggression and preceding development of malignancy. J Clin Oncol. 2004;22(14):2790–9.

51. Grasso CS, Wu YM, Robinson DR et al. The mutational landscape of lethal castration-resistant prostate cancer. Nature. 2012;487(7406):239-43.

52. Basu HS, Wilganowski N, Robertson S et al. Prostate cancer cells survive anti-androgen and mitochondrial metabolic inhibitors by modulating glycolysis and mitochondrial metabolic activities. Prostate. 2021;81(12):799–811.

53. Greene NP, Lee DE, Brown JL et al. Mitochondrial quality control, promoted by PGC-1alpha, is dysregulated by Western diet-induced obesity and partially restored by moderate physical activity in mice. Physiol Rep. 2015;3(7).

54. Solimando AG, Kalogirou C, Krebs M. Angiogenesis as Therapeutic Target in Metastatic Prostate Cancer - Narrowing the Gap Between Bench and Bedside. Front Immunol. 2022;13:842038.

55. Zielinski MR, Muenchow M, Wallig MA, Horn PL, Woods JA. Exercise delays allogeneic tumor growth and reduces intratumoral inflammation and vascularization. J Appl Physiol (1985). 2004;96(6):2249-56.

56. Norton KA, Popel AS. Effects of endothelial cell proliferation and migration rates in a computational model of sprouting angiogenesis. Sci Rep. 2016;6:36992.

57. McCullough DJ, Stabley JN, Siemann DW, Behnke BJ. Modulation of blood flow, hypoxia, and vascular function in orthotopic prostate tumors during exercise. J Natl Cancer Inst. 2014;106(4):dju036.

58. Warburg O, Wind F, Negelein E. The Metabolism of Tumors in the Body. J Gen Physiol. 1927;8(6):519–30.

59. Uo T, Sprenger CC, Plymate SR. Androgen Receptor Signaling and Metabolic and Cellular Plasticity During Progression to Castration Resistant Prostate Cancer. Front Oncol. 2020;10:580617.

60. Lin CS, Lee HT, Lee SY et al. High mitochondrial DNA copy number and bioenergetic function are associated with tumor invasion of esophageal squamous cell carcinoma cell lines. Int J Mol Sci. 2012;13(9):11228–46.

61. Caneba CA, Bellance N, Yang L, Pabst L, Nagrath D. Pyruvate uptake is increased in highly invasive ovarian cancer cells under anoikis conditions for anaplerosis, mitochondrial function, and migration. Am J Physiol Endocrinol Metab. 2012;303(8):E1036–52.

