## Supplemental Figure 1 for "Voluntary Exercise Attenuates Tumor Growth in a Preclinical Model of Castration-Resistant Prostate Cancer"

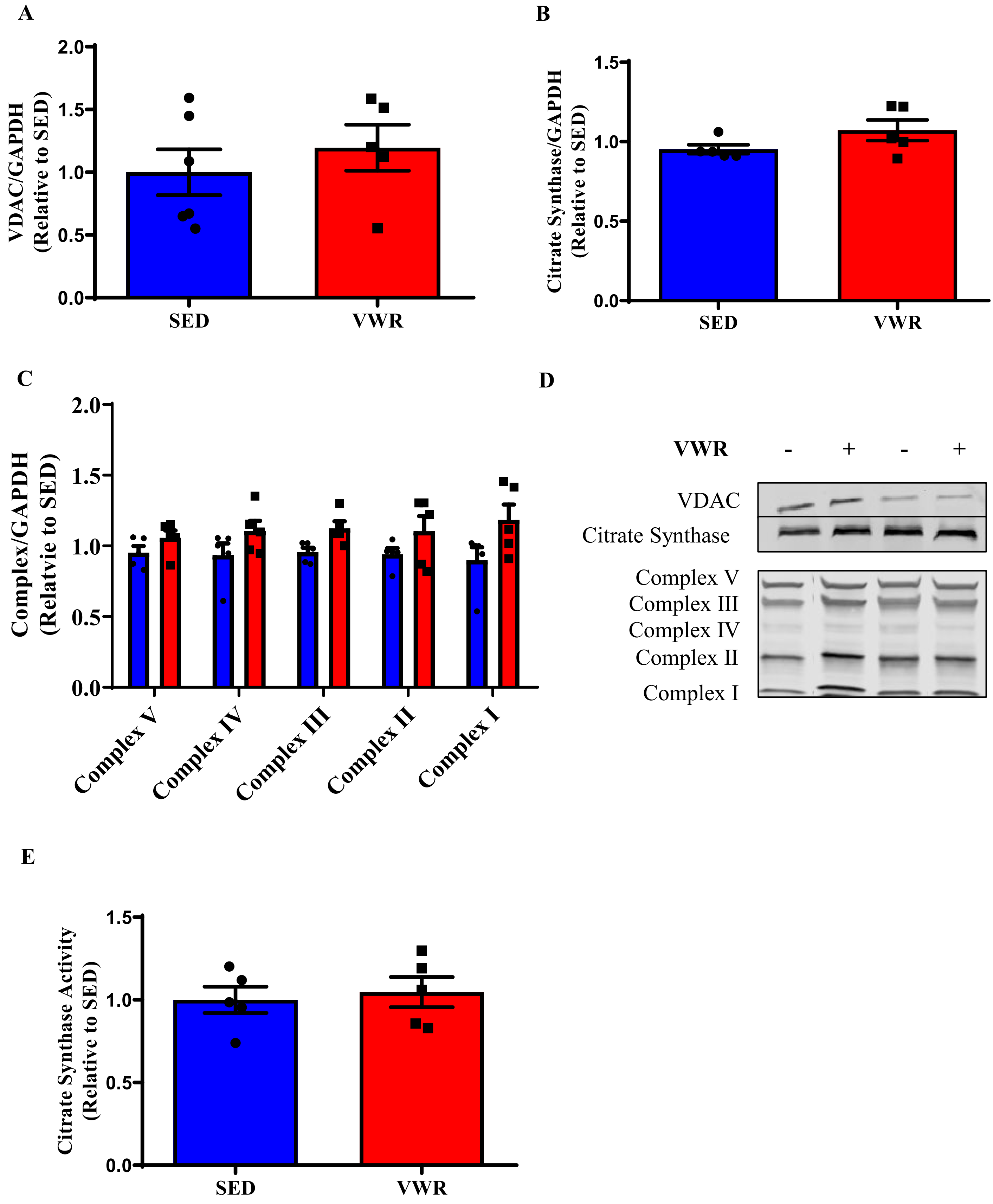

**Supplemental Figure 1. Voluntary wheel running did not change mitochondrial content in tumor tissue from a mouse model of CRPC.** (A) VDAC protein content. (B) Citrate synthase protein content. (C) Oxidative Phosphorylation Complex protein content. (D) Representative immunoblots for proteins in A-D. (E) Citrate synthase activity. N = 5-6/group. Data are expressed as Mean ± SEM.
