## Supplemental Table 1 for "Voluntary Exercise Attenuates Tumor Growth in a Preclinical Model of Castration-Resistant Prostate Cancer"

**Supplemental Table 1: Taqman probes used for the real-time quantitative polymerase chain reaction**

| **Gene Name** | **Assay Identifier** |
| --- | --- |
| *PSA/KLK3* | Hs02576345_m1 |
| *MCM2* | Hs01091564_m1 |
| *MCM6* | Hs00962418_m1 |
| *MCM7* | Hs00428518_m1 |
| *ELOVL5* | Hs01094704_m1 |
| *FKBP5* | Hs01561006_m1 |
| *DNM1L* | Hs01552605_g1 |
| *FIS1* | Hs01038322_m1 |
| *MFN1* | Hs00966851_m1 |
| *MFN2* | Hs00208382_m1 |
| *OPA1* | Hs01047013_m1 |
| *GAPDH* | Hs02786624_g1 |

Abbreviations: MFN1, Mitofusin 1; MFN2, Mitofusin 2; OPA1, Optic Atrophy 1; DNM1L, Dynamin-1-like protein; FIS1, Fission 1; GAPDH, glyceraldehyde-3 phosphate dehydrogenase.
