## Supplemental Table 2 for "Voluntary Exercise Attenuates Tumor Growth in a Preclinical Model of Castration-Resistant Prostate Cancer"

**Supplemental Table 2**: Complete list of antibodies

| **Antibody** | **Dilution** | **Supplier** | **Cat. No.** |
| --- | --- | --- | --- |
| Dynamin Related Protein 1 (Drp1) | 1:1000 | Cell Signaling Technology | 8570S |
| Fission 1 (FIS1) | 1:1000 | Santa Cruz Biotechnology | sc-376447 |
| Mitochondrial dynamic protein of 51kda (Mid51) | 1:1000 | Proteintech | 20164-1-AP |
| Mitofusin 2 (MFN2) | 1:1000 | Abcam | ab104274 |
| Optic Atrophy 1 (Opa1) | 1:1000 | Cell Signaling Technology | 67589S |
| P62 | 1:1000 | Abcam | ab56416 |
| Microtubule-associated proteins 1A/1B light chain 3B (LC3B) | 1:1000 | Cell Signaling Technology | 2775S |
| Citrate Synthase | 1:1000 | Abcam | ab96600 |
| Total OXPHOS Human Cocktail | 1:5000 | Abcam | ab110411 |
| Voltage-dependent anion channel (VDAC) | 1:1000 | Cell Signaling Technology | 4661S |
| Glyceraldehyde 3-phosphate dehydrogenase (GAPDH) | 1:5000 | Cell Signaling Technology | 2118S |
| IRDye 680RD Goat anti-Rabbit IgG Secondary Antibody | 1:5000 | LI-COR Biosciences | 926-68071 |
| IRDye 800RD Goat anti-Mouse IgG Secondary Antibody | 1:5000 | LI-COR Biosciences | 926-32210 |
