## Supplemental Table 4 for "Voluntary Exercise Attenuates Tumor Growth in a Preclinical Model of Castration-Resistant Prostate Cancer"

**Supplemental Table 4. Animal Characteristics**

|  | SED | VWR |
| --- | --- | --- |
| Body Weight Before Injection (g) | 26.9 ± 0.5 | 25.8 ± 0.5 |
| Final Body Weight (g) | 29.5 ± 0.9 | 29.0 ± 0.6 |
| Gastrocnemius Muscle Weight (g) | 0.156 ± 0.010 | 0.143 ± 0.008 |
| TA Muscle Weight (g) | 0.071 ± 0.010 | 0.069 ±0.007 |
| Heart Weight (g) | 0.137 ± 0.004 | 0.144 ± 0.007 |
| Epididymal Fat Pad Weight (g)  Daily Wheel Running Distance (meter) | 0.076 ± 0.005    n/a | 0.060 ± 0.003*  1547.7 ± 349.0 |

Data are presented as Mean ± SEM. *TA*, Tibialis Anterior. N=6/group. ***** P<0.05 vs. SED.
